# Preemptive minocycline decreases allodynia and depressive-like behaviors in a peripheral neuropathy rat model: a preliminary study

**DOI:** 10.1101/2024.10.23.619782

**Authors:** Anh Cong Tuan Le, Ângela Sousa, Nuno Dinis Alves, Hugo Leite-Almeida

## Abstract

**Background and Objective:** The neuroimmune system plays a critical role at all phases of chronic pain including at its onset. We therefore hypothesized that preemptive immunomodulation could decrease susceptibility and/or offer protection to pain.

**Materials and Methods:** Six days before the spared nerve injury (SNI), Wistar Han rats were treated with either the immunomodulator minocycline (MIN) or vehicle (VEH). After that, half of the animals from each group switch treatments for additional 7 days, resulting in 4 groups: continuous treatment (MIN/MIN), pretreatment (MIN/VEH), early treatment (VEH/MIN) and the control (VEH/VEH). Mechanical allodynia was recorded using Von Frey test until 4 weeks after SNI where depressive-like behaviors of the animals were also assessed using sucrose preference test.

**Results:** The continuous treatment provided sustained protection against mechanical allodynia, with rats in this group showing a significantly higher threshold to pain sensitivity compared to those in VEH/VEH. In contrast, pain relief effects were not observed with MIN/VEH and VEH/MIN. Additionally, animals in MIN/MIN, and VEH/MIN exhibited decreased anhedonic-like behavior at 30 days after SNI, relative to the control.

**Conclusions:** The exposure to an anti-inflammatory drug circa the installation of a neuropathic lesion had a positive impact on allodynia and on anhedonic behavior for a relatively long period after treatment cessation. The results support the assertions that pain trajectories can be altered at pain early stages by targeting the neuroimmune system. This proof-of-concept has the potential to be broadened to other drugs and/or therapeutical schemes.

**Highlights:** Chronic pain susceptibility varies across individuals.

(endo)phenotypes of susceptibility manifest prior/circa pain onset.

Preemptive minocycline treatment transiently decreases allodynia.

Preemptive minocycline treatment provides sustained reduction of depressive-like behavior.

Immune system modulation prior to pain onset or at early stages have beneficial outcomes.

## 1. INTRODUCTION

Pain is defined according to the International Association for the Study of Pain (IASP) as an unpleasant sensory and emotional experience associated with, or resembling that associated with, actual or potential tissue damage ^1^. Acute pain that persists or recurs for more than 3 months will be classified as chronic pain, which is often accompanied by comorbidities such as anxiety, depression, and sleep disturbances ^1,2^. Despite being broadly studied, the efficacy of pharmacological treatment for this condition is still inadequate ^3^. Developing new pharmacological approaches that improve the effectiveness is still an unmet demand.

Recent evidence in both humans and rodents suggests that chronic pain can be predicted from the early stages of the condition and even prior to the initial physiological incursions. For instance, preoperative anxiety, pain catastrophizing, and genetics in humans ^4–6^ or diet and exercise in rodents were all reported as predictors of chronic pain ^7–9^. Manipulating predictive or susceptibility factors to adjust the pain trajectory can be considered a novel method of chronic pain treatment.

Neuroimmune interaction is proved to contribute greatly the chronic pain pathogenesis ^10,11^, especially mediators released by immune cells like cytokines can initiate the acute-to-chronic pain transition. While attempts to correct the dysfunction or alter the interaction of these immune cells and neurons have widely been made to alleviate developed long-lasting pain ^12^, to our knowledge, so far there have not been studies investigate the possibility to prime pain pathways by modulating the neuroimmune system before pain onset. Minocycline (MIN) is a second-generation tetracycline known to suppress microglia activation - a CNS-located glia cell that is importantly involved in the mechanisms of neuropathic pain ^13–15^ . Animals treated with this compound showed substantially lower microglia-released cytokines and higher threshold to pain sensitivity in different models of neuropathy ^16–18^. Given that, in this preliminary study, we used an immunomodulator MIN with different preemptive regimes on spared nerve injury rats and assessed the effects on the following pain trajectories as well as the accompanying mood disorders^19^.

## 2. MATERIALS AND METHODS

### 2.1. Animals

Twenty-four 2-month-old male Wistar-Han rats were used in this experiment. Animals were pair-housed in standard cages with food and water available *ad libitum* and maintained in a room with controlled light (12-hour cycle; on 08:00) and temperature (22°C ± 1°C). Experiments were carried out by qualified FELASA category C researchers. All procedures involving animals were approved by the respective local organizations, and the experiments were performed according to the European Community Council Directive 2010/63/EU guidelines.

### 2.2. Experimental design

Animals were randomly distributed in 4 groups, 6 animals each. Treatments started 6 days prior to the neuropathy either with MIN or vehicle (VEH) (Figure 1A). After that, half of the animals in each group maintained the treatment for additional 7 days while the remaining animals crossed to the other group for the same amount of time resulting in 4 groups: continuous treatment (MIN/MIN)), pretreatment (MIN/VEH), early treatment (VEH/MIN) and the control (VEH/VEH). Mechanical allodynia was monitored for 4 weeks after SNI; anhedonic behavior was also assessed at the end of this period.

**Figure 1.**
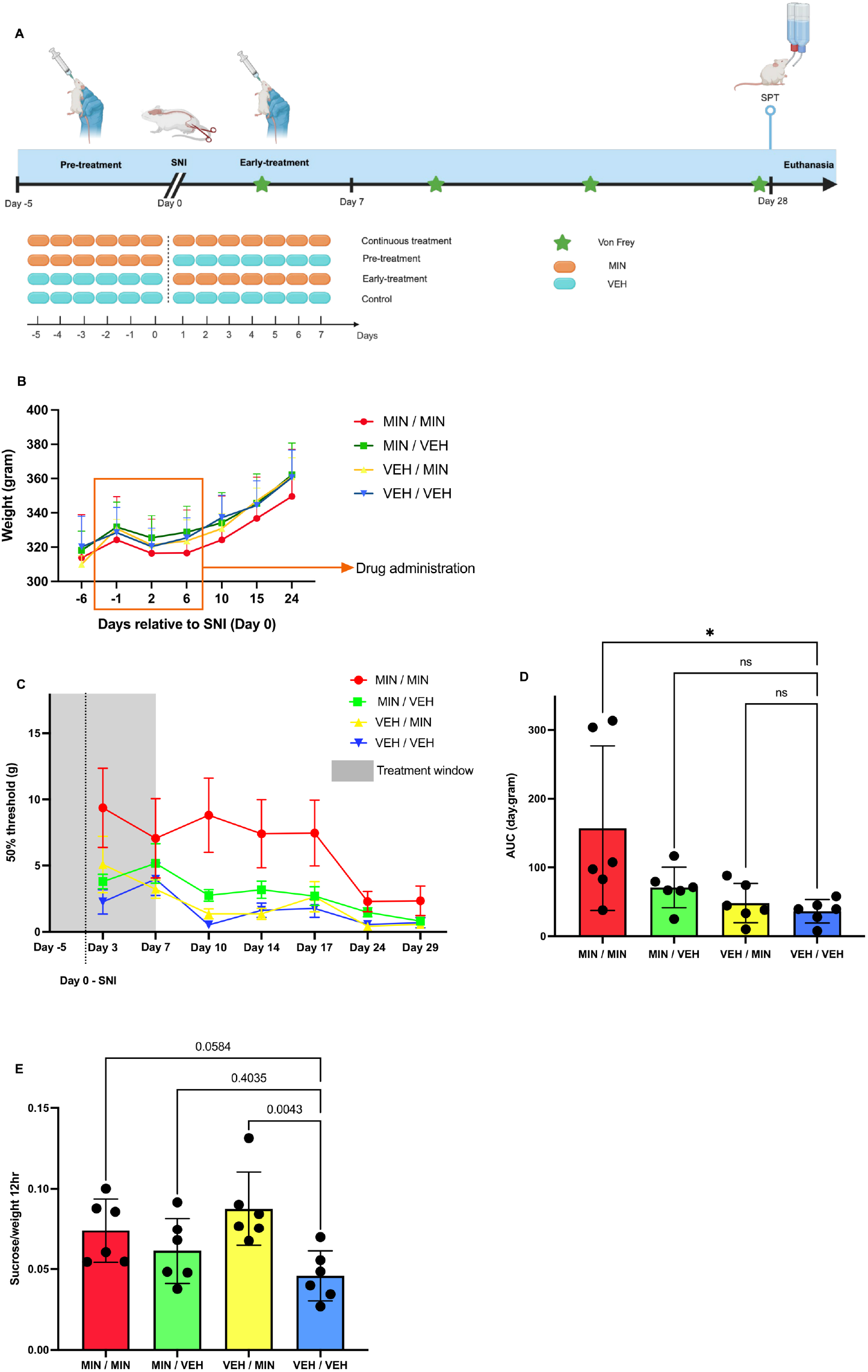
Effects of minocycline preemptive treatment on SNI rats. **(A)** Experimental timeline. Animals were handled for 1 week and then habituated to oral gavage for 4 days prior to the initiation of drug administration (not shown). Animals were then divided in 4 groups – MIN/MIN, MIN/VEH, VEH/MIN and VEH/VEH. At day 0, after the 6^*th*^ gavage, the SNI was implemented. **(B)** Weight evolution of animals preemptively treated with minocycline. Rats treated with minocycline showed no different in terms of weight evolution, compared to vehicle treated rats. **(C)** Effects of treatment on mechanical allodynia, presented overtime. **(D)** Cumulative effects of treatment on mechanical allodynia. Preemptive treatment with minocycline attenuated the severity of SNI-induced allodynia. Mechanical allodynia was assessed using the von Frey test, presented over time (C), and the cumulative effect AUC (D). **(E)** Effects of treatment on SPT. Depressive-like behavior was evaluated using SPT while the sucrose consumption per weight was used as index for comparisons. Preemptive treatment with minocycline attenuated SNI-induced anhedonia. ANOVA with repeated measure, one-way ANOVA, or two-way ANOVA were utilized where appropriate for statistical tests. Data was presented as mean ± SD, except for (C) where data was presented as mean ± SEM. n=6 male rats per group, *P<0.05, **P<0.01, ***P<0.001.

### 2.3. Drug administration

MIN (Minotrex, 100mg; Laboratório Medinfar, Amadora, Portugal), was diluted in sterile water and administered every day at 11 a.m., via oral gavage, using a dosage of 50mg/kg ^20,21^ . To avoid animals enduring stress during drug administration, they were carefully habituated to oral gavage with sterile water for 4 days before starting the experiments. Minocycline administration period was 13 days, starting 6 days (day -5) before the SNI and ending 7 days after (Figure 1A).

### 2.4. Neuropathy model

The spared nerve injury was performed as previously described by the group with minor alterations to the original description ^22–24^. Briefly, rats were anesthetized intraperitoneally 1ml/kg of a 1,5:1,0 mixture of ketamine (Imalgene, 100 mg/mL; Merial, Lyon, France) and medetomidine (Dormitor, 1 mg/mL; Orion Pharma, Espoo, Finland). An incision was made on the lateral surface of the right thigh through the biceps femoris muscle. Three sciatic nerve branches were exposed, and the tibial and common peroneal nerves were ligated. The section was performed distal to the ligation site, leaving the sural nerve intact. At the end of the procedure, the animals were administered with a subcutaneous injection of atipamezole hydrochloride (Antisedan®, Esteve Farma; 0.35 mg) ^25^ and monitored carefully for recovery.

### 2.5. Mechanical Allodynia

Mechanical allodynia was measured using von Frey (VF) test. Prior to the operation, animals were habituated to the apparatus and hypersensitivity was assessed with the up-and-down method as previously described by the group ^22,23^.

### 2.6. Sucrose Preference test

The Sucrose Preference test (SPT) was used as an indicative of anhedonia in the animals ^26,27^. The test was conducted on the 28^th^ day after SNI with a 4-day protocol. On the first day, at 9 a.m. two bottles including one with 300ml of a 1% sucrose solution and the other 300ml of water were placed to the animal cages. The next day at the same time the bottles switched sides to minimize any side preferences. On the third day, at 9 a.m. the rats were weighted and moved to the test room in a new cage, single-housed, and were put on food and water deprivation. Two bottles per cage were prepared, one containing 300ml of a 1% sucrose solution and the other 300ml of water. The bottles’ weight was recorded before randomly placing the bottles on the cages, and after 12 hours when the test finished, and the rats returned to their original cages and rooms. The sucrose consumption per animal weight was used as index for comparisons.

### 2.7. Statistical analysis

Statistical analyses were conducted in JASP version 0.18.1 and GraphPad Prism version 8.0.1, and Microsoft Office Excel. ANOVA with repeated measures followed by the Holm-Bonferroni method were performed to analyze mechanical allodynia. Additionally, the cumulative effect of treatments over time was determined as area under the curve (AUC). To analyze the results of the SPT test we performed one-way ANOVA with Dunnett, and independent t-test was also employed to give more insights about the data. Meanwhile two-way ANOVA with Tukey was used for examining weight evolution of animals. A p-value < 0,05 was considered statistically significant.

## 3. RESULTS

### 3.1. Minocycline exerted no significant impacts on animals’ weight evolution

Rats treated with minocycline showed no differences in terms of weight evolution, compared to vehicle treated rats (Figure 1B). A two-way ANOVA showed there was no effects on animals’ weights of Treatment [F (3,20) = 0.3819, p>0.05], nor Treatment x Time [F (18,20) = 1.504, p>0.05], but only effects of Time [F (3.344,66.88) = 180.4, p<0.0001], and Subjects [F (20,120) = 62.50, p<0.0001].

### 3.2. Preemptive minocycline treatment showed effects on mechanical allodynia

The threshold for mechanical stimulation in the control group (VEH/VEH) dropped from 15,003 ± 0 at baseline to 2,266 ± 0,926 on day 3 after SNI, which remained until day 29 (0,913 ± 0,562) (Figure 1C). ANOVA with repeated measures indicated an overall effect of minocycline Treatment [F (3,20) = 3,726, p = 0,030] and Time [F (7,126) = 65,038, p < 0,001] as well as a Treatment x Time interaction [F (21,126) = 1,752, p =0,031]. Although the following post-hoc analysis showed no further differences, when assessing the cumulative effects of treatments using the AUC, a one-way ANOVA revealed [F (3,20) = 4.392, p = 0,0157], with Dunnett’s multiple comparisons tests showed the MIN/MIN presented sustained significantly higher thresholds, compared to the control group (p< 0.05). Such effect was not present in the other 2 treatment groups (VEH/MIN and MIN/VEH) (Figure 1D).

### 3.3. Minocycline preemptive treatment significantly improved anhedonia

In terms of sucrose consumption, a one-way ANOVA revealed statistically significant differences between groups [F (3,20) = 4.882, p = 0.0105], with an effect size of 0.4227. Post-hoc analysis revealed that rats in the early-treatment VEH/MIN group were significantly less anhedonic, relative to VEH/VEH (p < 0.01), which the pretreatment failed to exhibit. Noticeably, the continuous treatment MIN/MIN effect showed a trend toward significance (p = 0.0584) (Figure 1E). Comparing directly to the VEH/VEH using independent t-test, this treatment exhibited better protection [t (10) = 2.741, p < 0.05].

## 4. DISCUSSION

Accumulating evidence shows that minocycline hampers microglia-driven sensitization and consequent pain ^13,14,28^. Our present study revealed that minocycline exposure around the neuropathic injury time improved pain trajectories in the observed period. This not only highlights the drug’s effects but also indicates that the period surrounding the pain onset is crucial, as modifying the immune system during this phase affects susceptibility to subsequent chronic pain.

The long-lasting effects of minocycline preemptive treatment found in our work are consistent with previous findings ^29,30^, where animals remained protected against hypersensitivity for up to 21 days. In comparison between regimes, MIN/MIN provided the best protection against mechanical allodynia, but as regards comorbid depression, not only MIN/MIN, VEH/MIN was also effective. The overall superiority of the continuous treatment aligns with observations from other studies, in which systematic microglia inhibitors are more effective when being administrated before injury and continue through the week after ^18,31^. Meanwhile, the attenuation of depressive-live behaviors suggests that preemptive minocycline impacts not only at the peripheral system but also centrally. The lack of pain relief results from VEH/MIN hinted there might exist different protective mechanisms against allodynia and depressive – like behaviors, which requires future works to confirm.

Another important aspect pertains with the time course of the effects. Minocycline protection lasted approximately 17 days, which goes well beyond the drug pharmacokinetics. This drug has an average half-life of 15.5 hours, meaning that 97% of the drug is eliminated from the plasma after 80 hours ^32,33^. In other words, it is cleaned from the system long before the 2–3-week allodynia protection period. A possible explanation for this is that minocycline’s main target is microglia and microglia activity changed overtime. Shortly after nerve injury, this immune cell enters its hyperactive state, which contributes greatly to the increase of hypersensitivity ^14^. After that period, the role of microglia becomes less significant as their activity is in the remission phase, which is accordance to the wear-off of the drug’s efficacy. Since factors contributing to chronic pain progression or resolution exist as a complex network including multiple cell types and pathways, a compound with broad spectrum effects might increase the protective period. It is still not clear why the protection effects on depressive-like behaviors lasts for a longer period.

In conclusion, our results support the assertions that (1) it is possible to prime the system prior to pain installation to change the pain trajectories, and (2) neuroimmune interaction is an important target for this purpose. This proof-of-concept can be expanded to other drugs and on both male and female subjects since mechanical allodynia is mediated by different component of the immune system ^16^.

## AUTHOR CONTRIBUTIONS

Anh Cong Tuan Le: Conceptualization, Methodology, Validation, Formal analysis, Investigation, Resources, Writing - Original Draft, Writing – Review & Editing, Visualization. Agreement to be accountable for all aspects of the work. Final approval of the version to be published.

Ângela Sousa: Methodology, Formal analysis, Investigation. Agreement to be accountable for all aspects of the work. Final approval of the version to be published.

Nuno Dinis Alves: Investigation. Writing – Review & Editing. Agreement to be accountable for all aspects of the work. Final approval of the version to be published.

Hugo Leite-Almeida: Conceptualization, Methodology, Validation, Resources, Writing – Review & Editing, Supervision, Project administration, Funding Acquisition. Agreement to be accountable for all aspects of the work. Final approval of the version to be published.

## STATEMENTS AND DECLARATIONS

### ETHICAL CONSIDERATIONS

All procedures involving animals were approved by the respective local organizations, and the experiments were performed according to the European Community Council Directive 2010/63/EU guidelines.

## DATA AVAILABILITY STATEMENT

Data associated with this study are presented in the article. Data files are available upon request.

